# Cell-specific CRISPR/Cas9 activation by microRNA-dependent expression of anti-CRISPR proteins

**DOI:** 10.1101/480384

**Authors:** Mareike D. Hoffmann, Sabine Aschenbrenner, Stefanie Grosse, Kleopatra Rapti, Claire Domenger, Julia Fakhiri, Manuel Mastel, Roland Eils, Dirk Grimm, Dominik Niopek

**Affiliations:** Synthetic Biology Group, Institute for Pharmacy and Biotechnology (IPMB) and Center for Quantitative Analysis of Molecular and Cellular Biosystems (BioQuant), University of Heidelberg, Heidelberg, 69120, Germany; Division of Theoretical Bioinformatics, German Cancer Research Center (DKFZ), Heidelberg, 69120, Germany; Department of Infectious Diseases, Virology, University Hospital Heidelberg, Heidelberg, 69120, Germany; BioQuant Center and Cluster of Excellence CellNetworks at Heidelberg University, Heidelberg, 69120, Germany; Digital Health Center, Berlin Institute of Health (BIH) and Charité, Berlin, 10178, Germany; Health Data Science Unit, University Hospital Heidelberg, Heidelberg, 69120, Germany; German Center for Infection Research (DZIF), partner site Heidelberg, Heidelberg, 69120, Germany

**Keywords:** CRISPR, anti-CRISPR protein, microRNA, Adeno-associated virus, synthetic biology

## Abstract

The rapid development of CRISPR/Cas technologies brought a personalized and targeted treatment of genetic disorders into closer reach. To render CRISPR-based therapies precise and safe, strategies to confine the activity of Cas(9) to selected cells and tissues are highly desired. Here, we developed a cell type-specific Cas-ON switch based on miRNA-regulated expression of anti-CRISPR (Acr) proteins. We inserted target sites for miR-122 or miR-1, which are abundant specifically in liver and muscle cells, respectively, into the 3’UTR of Acr transgenes. Co-expressing these with Cas9 and sgRNAs resulted in Acr knockdown and correspondingly in Cas9 activation solely in hepatocytes or myocytes, while Cas9 was efficiently inhibited in off-target cells. We demonstrate control of genome editing and gene activation using a miR-dependent AcrIIA4 in combination with different *Streptococcus pyogenes (*S*py)*Cas9 variants (full-length Cas9, split-Cas9, dCas9-VP64). Finally, to showcase its modularity, we adapted our Cas-ON system to the smaller and more target-specific *Neisseria meningitidis (Nme)*Cas9 orthologue and its cognate inhibitors AcrIIC1 and AcrIIC3. Our Cas-ON switch should facilitate cell-specific activation of any CRISPR/Cas orthologue, for which a potent anti-CRISPR protein is known.

## INTRODUCTION

CRISPR (clustered regularly-interspaced short palindromic repeats) technologies provide an efficient and simple means to perform targeted genetic manipulations in living cells and animals (1–5). Today, the rapidly expanding CRISPR toolbox enables genomic knock-ins/outs (2,3), gene silencing and activation (6–8), epigenetic reprogramming (9–11), as well as single-base editing (12,13). These tools facilitate detailed genetic studies in cells and animals and hold enormous potential for the future treatment of genetic disorders (14).

With respect to *in vivo* application of CRISPR technologies, strategies to confine CRISPR/Cas9 activity to selected cells and tissues are highly desired. For genetic studies in animals, for instance, confining perturbations to selected cells is critical when aiming at disentangling the role of selected cell types in a particular phenotype or simply to avoid negative side-effects and/or artefacts that would arise from unspecific perturbations. Moreover, in the context of therapeutic genome editing within human patients, ensuring maximum specificity and hence safety of a treatment is absolutely critical. Until today, however, virtually any mode of efficient *in vivo* delivery of the CRISPR/Cas components (e.g. via viral vectors, nanoparticles, lipophilic complexes etc.) is likely to affect many cell types and tissues beyond the one of actual (therapeutic) interest. This limited specificity, in turn, causes substantial risks of (treatment) side-effects (14,15).

One strategy to address this limitation would be to render the activity of the CRISPR components dependent on endogenous, cell-specific signals, so that the genetic perturbation is induced solely in the target cell population, but not in off-target cells. One such signal are mi(cro)RNAs, i.e. small, regulatory and non-coding RNAs that are involved in eukaryotic gene expression control (16,17). Being part of the RNA-induced silencing complex (RISC), miRNAs recognize sequence motifs present on m(essenger)RNAs that are complementary to the miRNA sequence. The RISC then typically mediates mRNA degradation, or translation inhibition or both, thereby causing a gene expression knockdown (16,17).

More than 1000 miRNAs have been described in humans (http://www.mirbase.org), and many miRNAs or miRNA combinations have been identified, which occur exclusively in selected cell types or disease states (18–23). These include, for instance, miR-122, which is selectively expressed in hepatocytes (18), or miR-1, which is highly abundant in myocytes (22,23). Such unique signatures have in the past been successfully harnessed for cell-specific expression of transgenes in cultured cells and mice (24,25). Adapting this strategy to CRISPR/Cas would thus offer an effective means to confine CRISPR-mediated perturbations to selected cell types.

We have previously shown that integrating miRNA-122 binding sites into the 3’UTR (3’ untranslated region) of a CRISPR/Cas9 transgene can be used to de-target Cas9 expression from hepatocytes (26). A subsequent study by Hirohide Saito’s group expanded this approach to further miRNA candidates (miR-21 and miR-302a) (27). Moreover, they added a negative feedback loop to the system, thereby establishing a positive relation between miRNA abundance and Cas9 activity (27). To this end, the authors expressed Cas9 from an mRNA harbouring an L7Ae binding motif (K-turn), while co-expressing the L7Ae repressor from an mRNA carrying miRNA binding sites in its 3’UTR (27). The resulting Cas-ON switch enabled miRNA-dependent Cas9 activation. The system was leaky, however, and showed a less than 2-fold dynamic range of regulation, thereby limiting its utility for *in vivo* applications.

Here, we created a novel, robust and highly flexible cell type-specific Cas9-ON switch based on anti-CRISPR proteins (28–32) expressed from miRNA-dependent vectors. We placed AcrIIA4, a recently discovered *Streptococcus pyogenes (Spy)*Cas9 inhibitor, under miR-122 and miR-1 control, thereby enabling hepatocyte- or myocyte-specific activation of various *(Spy)*Cas9 variants (full-length Cas9, split-Cas9, dCas9-VP64) with an up to ∼100-fold dynamic range of regulation. Finally, to demonstrate its modularity, we expanded our Cas-ON approach to the smaller and more target-specific *Neisseria meningitidis* (*Nme*)Cas9 and its cognate inhibitors AcrIIC1 and AcrIIC3.

## MATERIAL AND METHODS

### Cloning

A list of all constructs used and created in this study is shown in Supplementary Table 1. Annotated vector sequences are provided as Supplementary Data (GenBank files). Plasmids were created using classical restriction enzyme cloning, Golden Gate Assembly (33) or Gibson assembly (New England Biolabs). Oligonucleotides were obtained from Integrated DNA Technologies (IDT) or Sigma-Aldrich. Synthetic, double-stranded DNA fragments (gBlocks) were obtained from IDT.

Luciferase knockdown reporters carrying miRNA binding sites within the 3’UTR of the *Renilla* luciferase gene (pSiCheck-2 no miR target site/ 2xmiR-122 target sites) were generated by inserting a DNA fragment encoding two miRNA target sites followed by a bovine growth hormone (BGH) polyA signal into the psiCheck2 vector (Promega) via XhoI/NotI. The CMV promoter-driven *Spy*Cas9 expression vector (Addgene plasmid # 113033) was previously developed by us (34). A *Spy*Cas9-GFP fusion was cloned by PCR-amplifying EGFP from vector pSpCas9(BB)-2A-GFP (Addgene plasmid# 48138, which was a kind gift from Feng Zhang) followed by insertion of the PCR amplicon into the *Spy*Cas9 vector via EcoRI/HindIII. The AcrIIA4 coding sequence and the mCherry-AcrIIA4 coding sequence were obtained as human codon-optimized, synthetic DNA fragments from IDT and cloned into pcDNA3.1^(-)^ (ThermoFisher) via NheI/NotI. 2xmiR-122 target sites or a scaffold sequence identical in length but lacking the miR target sites were inserted into the resulting vectors by oligo cloning via EcoRI/HindIII, yielding vectors CMV-(mCherry)-AcrIIA4–2x*miR-122* and CMV-(mCherry)-AcrIIA4–*scaffold*.

The luciferase cleavage reporter for measuring *Spy*Cas9 activity was previously reported by us (34). It comprises an SV40 promoter-driven *Renilla* luciferase gene, a TK promotor-driven Firefly luciferase gene, and an H1 promoter-driven sgRNA targeting the Firefly luciferase gene. The pRL-TK vector encoding *Renilla* luciferase was obtained from Promega. AAV vectors encoding (i) *Spy*Cas9 (Addgene #113034) or (ii) an H1 or U6 promoter-driven sgRNA (F+E scaffold (35)) and a RSV promoter-driven EGFP (Addgene #113039) were previously reported by us (36). Annealed oligonucleotides corresponding to the genomic target site were cloned into the sgRNA AAV vector via BbsI using Golden Gate cloning (33). All sgRNA target sites relevant to this study are shown in Supplementary Table 2. AAV vectors encoding CMV or EF1α promoter-driven AcrIIA4 variants were created by replacing the RSV promoter-driven GFP expression cassette from the sgRNA plasmids (36) with synthetic DNA fragments encoding CMV-FLAG-AcrIIA4-*scaffold*, CMV-FLAG-AcrIIA4*-2xmiR122*, CMV-FLAG-AcrIIA4*-2xmiR1*, EF1α-AcrIIA4-*scaffold* or EF1α-AcrIIA4-*2xmiR122*, all succeeded by a BGH terminator sequence. An AAV vector co-encoding an N-terminal *Spy*Cas9 fragment fused to a split-intein and a U6 promoter-driven sgRNA scaffold (F+E) was generated by inserting a DNA fragment encoding the U6-promoter-sgRNA scaffold via MluI/XbaI into vector pAAV-SMVP-Cas9N (kind gift from George Church (Addgene plasmid # 80930)). An AAV vector co-encoding the corresponding C-terminal *Spy*Cas9 fragment fused to a split-intein was a kind gift from George Church (Addgene plasmid # 80931). CMV promoter-driven AcrIIA4 fragments with or without 2xmiR-122 target sites were introduced into this vector by first inserting unique XbaI and MluI sites behind the SV40 polyA. These were subsequently used to introduce CMV-AcrIIA4-*scaffold* and CMV-AcrIIA4-*2xmiR-122* fragments generated by PCR from corresponding template vectors described above.

The pAAV-pSi vector co-encoding Firefly and *Renilla* luciferase (pAAV-pSi) was previously reported by us (36). A single miR-1 binding site was introduced into the *Renilla* luciferase gene 3’UTR by oligo cloning via XhoI/NotI resulting in the vector pAAV-pSi 1xmiR target site. The Tet-inducible luciferase reporter and corresponding sgRNA construct (sgRNA1_Tet-inducible Luciferase reporter) were kind gifts from Moritoshi Sato (Addgene plasmids # 64127 and # 64161). dCas9-VP64_GFP was a kind gift from Feng Zhang (Addgene plasmid # 61422). The pEJS654 All-in-One AAV-sgRNA-hNmeCas9 vector was a kind gift from Erik Sontheimer (Addgene plasmid # 112139). The VEGFA target site (NTS-33; Amrani *et al*., 2018, BioRxiv: https://doi.org/10.1101/172650) was introduced into this vector by oligo cloning via SapI. The AcrIIC1 and AcrIIC3 coding sequences were obtained as human codon-optimized, synthetic DNA fragment from IDT and cloned into the AAV CMV-driven AcrIIA4*-scaffold or* AAV CMV-driven AcrIIA4*-2xmiR122* vector by replacing the AcrIIA4 sequence with the AcrIIC1 or AcrIIC3 coding sequences.

In all cloning procedures, PCRs were performed using Phusion Flash High-Fidelity polymerase (ThermoFisher) or Q5 Hot Start High-Fidelity DNA Polymerase (New England Biolabs) followed by agarose gel electrophoresis to analyse PCR products. Bands of the expected size were cut out and the DNA was extracted by using a QIAquick Gel Extraction Kit (Qiagen). Restriction digests and ligations were performed with enzymes from New England Biolabs and ThermoFisher and according to the manufacturer’s protocols. Chemically-competent Top10 cells (ThermoFisher) were used for plasmid amplification and plasmid DNA was purified using the QIAamp DNA Mini, Plasmid Plus Midi or Plasmid Maxi Kit (all from Qiagen).

### Cell culture

Cells lines were cultured at 5 % CO_2_ and 37 °C in a humidified incubator and passaged when reaching 70 to 90 % confluency. HeLa and HEK293T cells were maintained in 1x DMEM without phenol red (ThermoFisher) supplemented with 10 % (v/v) fetal calf serum (Biochrom AG), 2 mM L-glutamine and 100 U per ml penicillin/100 µg per ml streptomycin (both ThermoFisher). Huh-7 medium was additionally supplemented with 1 mM non-essential amino acids (ThermoFisher). HeLa, HEK293T and Huh-7 cells were authenticated and tested for mycoplasma contamination prior to use via a commercial service (Multiplexion). HL-1 cells were cultured in Claycomb medium supplemented with 10 % (v/v) fetal calf serum, 1 % penicillin/streptomycin, 0.1 mM norepinephrine, and 2 mM L-Glutamine on plates pre-coated with 0.02 % (w/v) gelatin and 5 μg per ml fibronectin (all Sigma-Aldrich).

Cells were transfected with Lipofectamine^™^2000, Lipofectamine^™^3000 (both ThermoFisher) or jetPRIME^®^ (Polyplus-transfection) according to the manufacturers’ protocols and as specified in the experimental sections below.

### Fluorescence microscopy

HeLa and Huh-7 cells were seeded into 8-well Glass Bottom µ-Slides (ibidi) at a density of 30,000 cells per well for HeLa cells and of 60,000 cells per well for Huh-7 cells in 300 µl media. The next day, cells were transfected using 1.85 µl Lipofectamine^™^2000 per well by following the manufacturer’s protocol. The total amount of transfected DNA per well was 720 ng evenly split between plasmids encoding *Spy*Cas9-GFP, either mCherry-AcrIIA4*-scaffold* or AcrIIA4-*2xmiR-122*, and the luciferase cleavage reporter plasmid (to provide a sgRNA and an exogenous target). The medium was exchanged 6 h post transfection.

24 h post-transfection, HeLa and Huh-7 cells were treated with Hoechst 33342 solution at a final concentration of 5 µg per ml for 15 min at 37 °C. Then, the medium was replaced and imaging was performed using a Leica SP8 confocal laser scanning microscope equipped with automated CO_2_ and temperature control, a UV, argon, and a solid state laser, as well as a HCX PL APO 20x oil objective (N/A = 0.7). mCherry fluorescence was recorded using the 552 nm laser line for excitation and the detection wavelength was set to 578-789 nm. GFP fluorescence was recorded using the 488 nm laser line for excitation and the detection wavelength was set to 493-578 nm.

### Western blot

For Western blot analysis, cells were seeded into 6-well plates (CytoOne) at a density of 300,000 cells per well for HeLa cells and 450,000 cells per well for Huh-7 cells. The following day, cells were co-transfected with 500 ng of *Spy*Cas9-GFP and either 500 ng CMV-mCherry-AcrIIA4*-scaffold* or CMV-mCherry-AcrIIA4*-2xmiR-122* per well using Lipofectamine^™^3000. 24 h post-transfection, the media was aspirated from the culture plates and the cells were washed with ice-cold PBS. 50 μl of protein lysis buffer (150 mM NaCl, 10 mM TRIS, 1 mM EDTA, 0.5 % NP 40, and 10 % cOmplete Protease Inhibitor (Roche)) were added, and the cells were detached from the culture plate surface using a cell scraper. The cell suspension was then transferred into a 1.5 ml tube, incubated for 20 min on ice, and centrifuged for 10 min at 13,200 rpm (15,974 xg) and 4 °C. The supernatant containing the protein lysate was transferred into a new 1.5 ml tube and protein concentrations were measured using the Bradford Reagent (Sigma-Aldrich) according to the manufacturer’s protocol. 50 μg of protein lysate were then mixed with 4x Laemmli Sample Buffer (BIO-RAD), and the volume of each sample was adjusted to 30 µl using lysis buffer. The samples were denatured for 10 min at 95 °C, cooled down on ice and loaded on a 10 % Bis TRIS gel (Life Technologies). Proteins were then separated by molecular weight by applying 130 V for 120 min in 1x MOPS buffer (Life Technologies). Next, proteins were transferred onto a nitrocellulose membrane (pore size 0.2 μm) (Millipore) by using 1x borate transfer buffer (20 mM boric acid, 1 mM EDTA, 6.25 mM NaOH) and applying 300 mA current over night. The membrane was then cut at ∼72 kDa and ∼45 kDa, and washed in TBS-T (Tris-buffered saline (20 mM TRIS, 140 mM NaCl, pH7.6) supplemented with 0.1 % Tween 20 (Sigma-Aldrich)) and blocked by incubation in 5 % milk (skim milk powder, GERBU Biotechnik GmbH, diluted in TBS-T) at room temperature for 1 h. GFP antibody (ChromoTek, diluted 1:1,500) was used for *Spy*Cas9-GFP detection (190 kDa), α-tubulin antibody (Santa Cruz Biotechnology, diluted 1:500) was used for α-tubulin detection (55 kDa) and RFP antibody (ChromoTek, diluted 1:500) was used for mCherry-AcrIIA4 detection (38 kDa). All antibodies were diluted in 5 % milk in TBS-T, added to the corresponding membrane piece and incubated overnight at 4 °C while shaking. The next day, the membrane was washed three times for 5 min in TBS-T followed by incubation with HRP-(horse radish peroxidase-)linked secondary antibodies (anti-mouse antibody, 1:5,000 in 5 % milk in TBS-T (Dianova) or anti-rat antibody, 1:1,000 in 5 % milk in TBS-T (Jackson ImmunoResearch)) for 1 h at room temperature. The membrane was then washed three times for 5 min in TBS-T to remove unbound antibodies and SuperSignal™ West Pico PLUS Chemiluminescent Substrate (ThermoFisher) was applied for 5 min. Finally, the luminescence signal was detected using a ChemoStar detector (INTAS). Full-length Western blot images are shown in Supplementary Figure 1.

**Figure 1.**
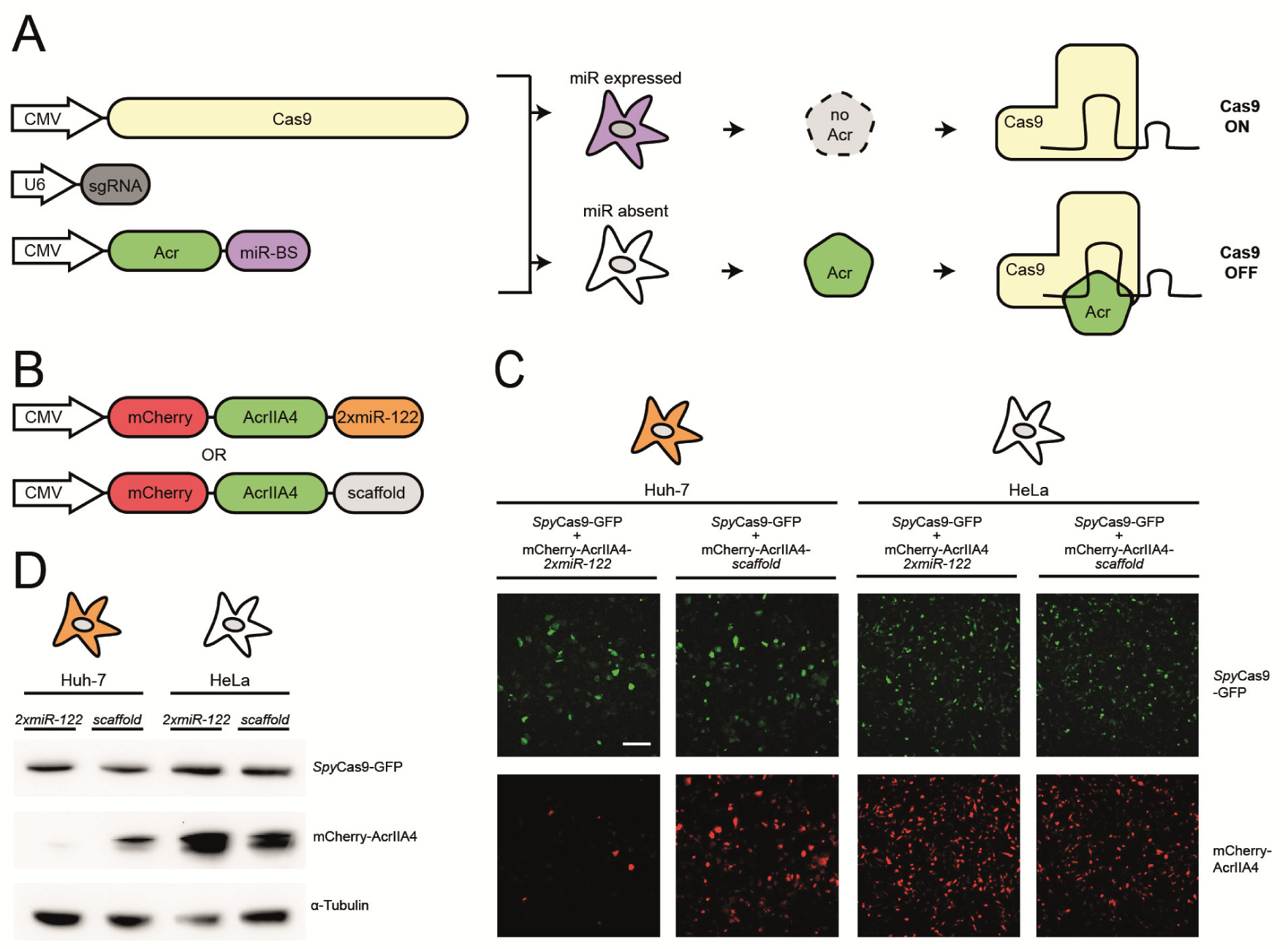
Designing a cell-specific Cas-ON switch based on miRNA-regulated anti-CRISPR genes. (**A**) Schematic of the Cas-ON switch. Target sites for cell-specific, abundant miRNAs are inserted into the 3’UTR of an Acr-encoding transgene. Upon delivery, a knockdown of Acr expression occurs selectively within the target cells, resulting in CRISPR/Cas activation. In OFF-target cells lacking the miRNA signature, the Acr protein is translated and inhibits CRISPR/Cas. (**B**) Schematics of mCherry-AcrIIA4 fusion constructs with or without 2xmiR-122 target sites. (**C, D**) Hepatocyte-specific knockdown of mCherry-AcrIIA4 expression. Huh-7 and HeLa cells co-transfected with *Spy*Cas9-GFP and either mCherry-AcrIIA4*-scaffold* or mCherry-AcrIIA4*-2xmiR-122*. (**C**) Representative fluorescence images from n = 2 independent experiments. Scale bar is 200 µm. (**D**) Complementary Western blot analysis of *Spy*Cas9-GFP and mCherry-AcrIIA4 expression. Data represent a single experiment.

### Luciferase assays

For luciferase experiments, HeLa and Huh-7 cells were seeded at a density of 6,000 cells per well, HEK293T cells were seeded at a density of 12,500 cells per well, and HL-1 cells were seeded at a density of 12,000 cells per well into 96-well plates (Eppendorf) using 100 µl culture medium per well. 16 h post-seeding, cells were either transiently transfected or transduced with AAV vectors as specified below (all plasmid/vector amounts are per well).

For miR-122- or miR-1-induced *Renilla* luciferase knockdown experiments (Supplementary Figures 3 and 7), Huh-7, HeLa and HEK293T cells were transfected with 20 ng of pSiCheck-2 reporter (with or without 2xmiR-122 target sites within the *Renilla* luciferase 3’UTR) and 80 ng of an irrelevant stuffer DNA (pcDNA3.1^(-)^, ThermoFisher). HL-1 cells were pre-treated with 0.5 μM of Bortezomib (Biomol) to improve transduction efficiency and then transduced with 10 µl pAAV-pSi vector per well (with or without miR-1 binding site within the *Renilla* 3’UTR; see below for AAV production).

For *Spy*Cas9 luciferase cleavage experiments (Figures 2A and Supplementary Figure 4), cells were co-transfected with 20 ng luciferase cleavage reporter plasmid, 20 ng CMV-*Spy*Cas9 expression vector, and different doses of AcrIIA4-*scaffold* or AcrIIA4*-2xmiR122* expression vectors (0.25, 1, 5 or 20 ng). For split-*Spy*Cas9 luciferase cleavage experiments (Supplementary Figure 6), cells were co-transfected with 20 ng luciferase cleavage reporter plasmid, 20 ng of AAV N-*Spy*Cas9-Intein and 20 ng of either (i) AAV Intein-C-*Spy*Cas9, (ii) AAV Intein-C-*Spy*Cas9-CMV-AcrIIA4*-scaffold* or (iii) AAV Intein-C-*Spy*Cas9-CMV-AcrIIA4*-2xmiR122*. To keep the total amount of DNA transfected constant between all samples, DNA was topped up to 100 ng per well using an irrelevant stuffer DNA (pcDNA3.1^(-)^).

**Figure 2.**
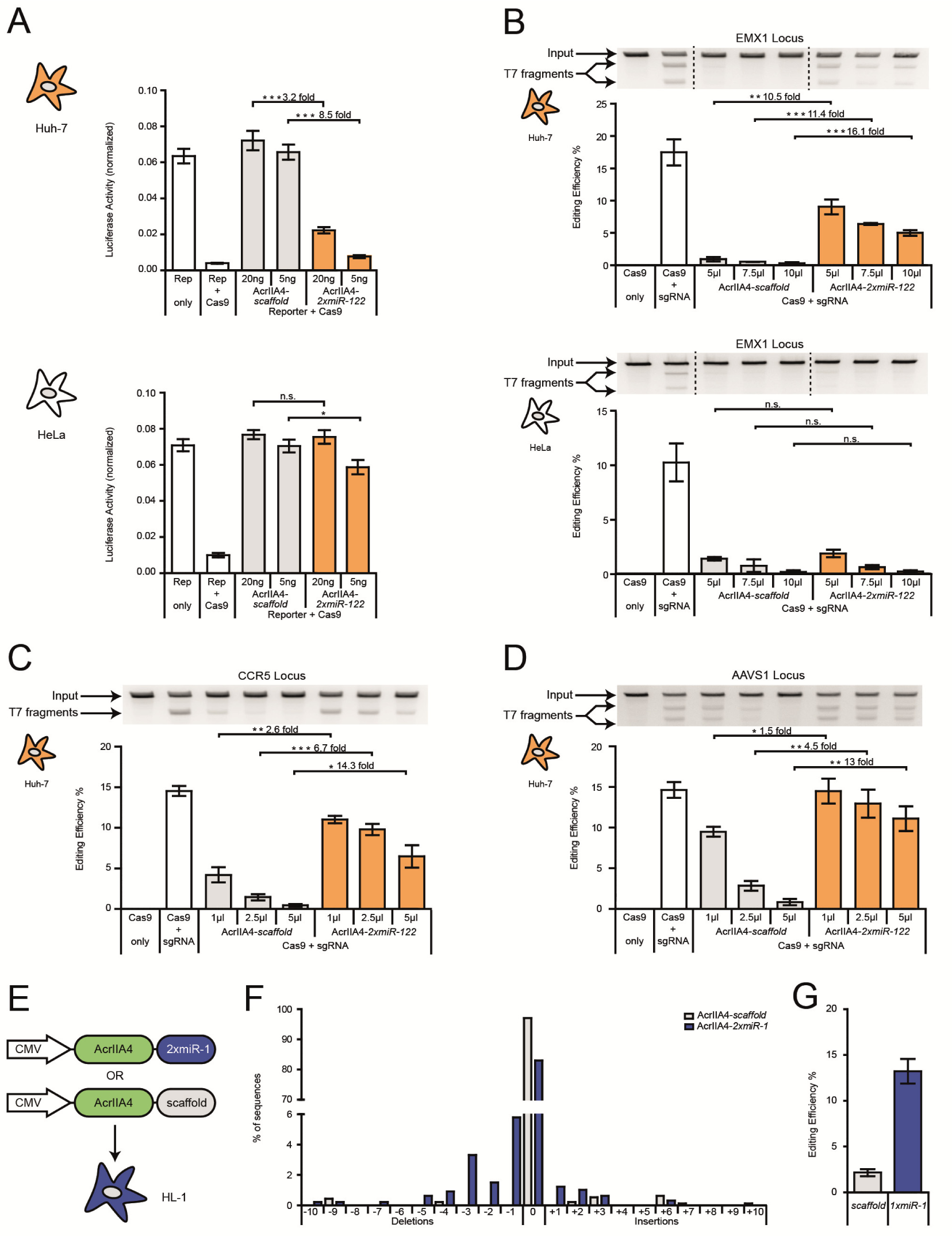
Hepatocyte- or myocyte-specific genome editing. (**A**) Hepatocyte-specific luciferase reporter cleavage via miR-122-dependent *Spy*Cas9 activation. Huh-7 or HeLa cells were co-transfected with plasmids encoding *Spy*Cas9, a luciferase reporter, a reporter-targeting sgRNA, and AcrIIA4*-miR-122* or AcrIIA4*-scaffold*, followed by luciferase assay. During transfection, the Acr vector dose was varied as indicated. Data are means ± s.e.m. (n = 7 independent experiments). (**B**) Hepatocyte-specific indel mutation of the human EMX1 locus. Huh-7 and HeLa cells were co-transduced with AAV vectors encoding *Spy*Cas9, an EMX-1-targeting sgRNA, and the indicated AcrIIA4 variant, followed by T7 endonuclease assay. During transduction, the Acr vector dose was varied as indicated. Data are means ± s.e.m. (n = 3 independent experiments). (**C, D**) Huh-7 cells were co-transduced with AAV vectors encoding *Spy*Cas9, a sgRNA targeting the human CCR5 (**C**) or AAVS1 (**D**) locus and the indicated AcrIIA4 variant, followed by T7 endonuclease assay. During transduction, the Acr vector dose was varied as indicated. Data are means ± s.e.m. (n = 3 independent experiments). (**A-D**) n.s. = not significant,\**P* < 0.05, \*\**P* < 0.01, \*\*\**P* < 0.001, by two-sided Student’s *t*-test. (**E**) Schematic of AcrIIA4 vectors for myocyte-specific genome editing. (**F-G**) MiR-1-dependent indel mutation of the Rosa-26 locus in myocytes. HL-1 cells were co-transduced with AAV vectors expressing *Spy*Cas9, a sgRNA targeting the murine Rosa-26 locus, and AcrIIA4 either with or without miR-1 binding sites, followed by TIDE sequencing. (**F**) Detailed analysis of indels observed at the Rosa-26 locus. Data for a representative sample is shown. (**G**) Quantification of the overall editing efficiency. Data are means ± s.e.m. (n = 3 independent experiments).

**Figure 3.**
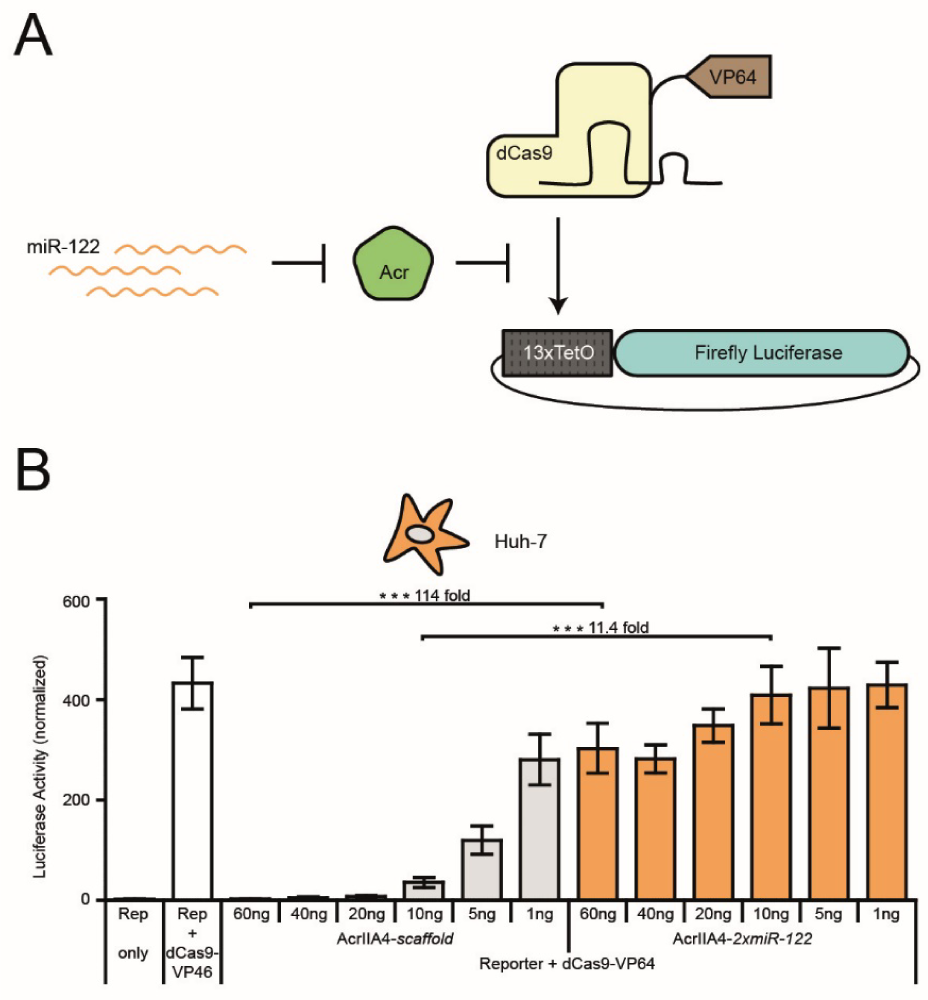
MiR-122-dependent gene activation via *Spy*dCas9-VP64. (**A**) Schematic of miR-122-dependent activation of luciferase reporter expression. MiR-122-dependent knockdown of AcrIIA4 releases *Spy*dCas9-VP64 activity and results in activation of luciferase expression. (**B**) Huh-7 cells were co-transfected with vectors encoding *Spy*dCas9-VP64, a luciferase reporter driven by a minimal promoter preceded by tet operator (TetO) sites, a TetO-targeting sgRNA and AcrIIA4*-miR-122* or AcrIIA4*-scaffold* construct (as control), followed by a luciferase assay. During transduction, the Acr vector dose was varied as indicated. Data are means ± s.e.m. (n = 4 independent experiments). \*\*\**P* < 0.001, by two-sided Student’s *t*-test.

**Figure 4.**
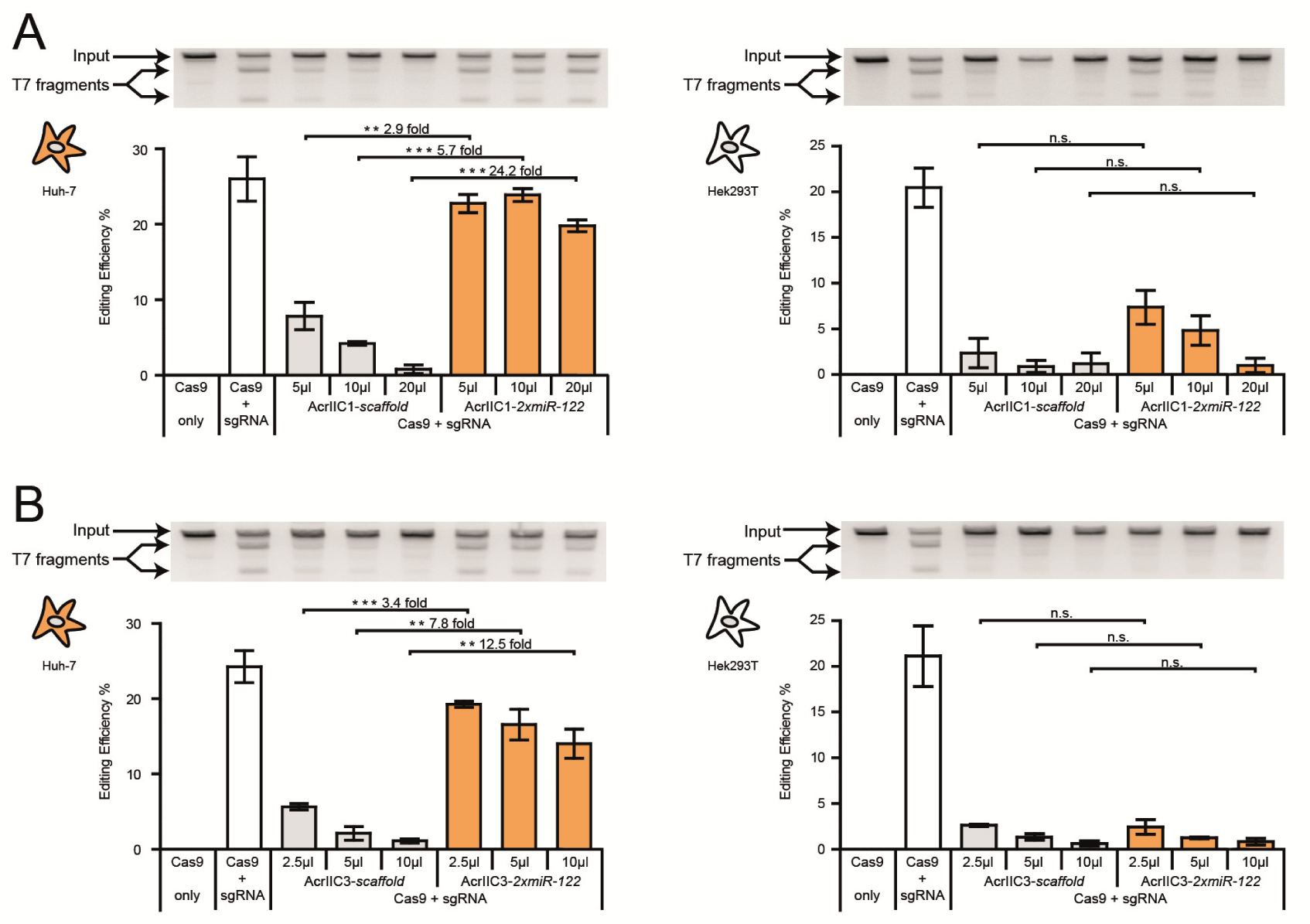
Hepatocyte-specific activation of *Nme*Cas9. (**A, B**) Huh-7 or HEK293T cells were co-transduced with AAV vectors expressing *Nme*Cas9, a sgRNA targeting the human VEGFA locus and either AcrIIC1 (**A**) or AcrIIC3 (**B**) carrying two miR-122 target sites or not, followed by T7 endonuclease assay. During transduction, the Acr vector dose was varied as indicated. Data are means ± s.e.m. (n = 3 independent experiments). n.s. = not significant, \*\**P* < 0.01, \*\*\**P* < 0.001, by two-sided Student’s *t*-test.

For *Spy*dCas9-VP64-mediated luciferase activation experiments (Figure 3B), cells were co-transfected with 20 ng Tet-inducible luciferase reporter plasmid, 20 ng dCAS9-VP64_GFP expression vector, 20 ng sgRNA1_Tet-inducible luciferase reporter, 1 ng pRL-TK (encoding *Renilla* luciferase, Promega), and different doses (1, 5, 10, 20, 40 or 60 ng) AcrIIA4 expression vector. DNA was topped up to 101 ng per well using an irrelevant stuffer DNA (pcDNA3.1^(-)^).

6 h post-transfection or 4 h post-transduction, the medium was replaced. HeLa, Huh-7 and HEK293T cells were incubated for 48 h and HL-1 cells were incubated for 72 h before measuring *Renilla* and Firefly luciferase activity using a Dual-Glo Luciferase Assay System (Promega) according to the manufacturer’s recommendation. In brief, cells were harvested in the supplied lysis buffer and Firefly and *Renilla* luciferase activities were measured using a GLOMAX^™^ Discover or GLOMAX^™^ 96-microplate luminometer (both Promega). Integration time was 10 s, and delay time between automated substrate injection and measurement was 2 s. For the miRNA-dependent luciferase knockdown experiments based on the pSiCheck-2 and pAAV-pSi vectors, *Renilla* luciferase photon counts were normalized to the Firefly luciferase photon counts (as miR target sites were inserted into the *Renilla* luciferase 3’UTR). For *Spy*Cas9 luciferase cleavage experiments and *Spy*dCas9-VP64-mediated luciferase activation experiments, Firefly luciferase photon counts were normalized to *Renilla* photon counts (as Cas9 is targeting the Firefly luciferase gene in these assays).

### AAV lysate production

For production of AAV-containing cell lysates, low-passage HEK293T cells were seeded into 6-well plates (CytoOne) at a density of 350,000 cells per well. The following day, cells were triple-transfected with (i) the AAV vector plasmid, (ii) an AAV helper plasmid carrying AAV serotype 2 (AAV2) *rep* and either the AAV2 (when targeting Huh-7, HeLa or HEK293Tcells) or AAV6 *cap* genes (when targeting HL-1 cells), and (iii) an adenoviral plasmid providing helper functions for AAV production, using 1.33 µg of each construct and 8 µl of TurboFect Transfection Reagent (ThermoFisher) per well. The AAV vector plasmid encoded either (i) *Spy*Cas9 driven from an engineered, short CMV promoter, (ii) a U6 promoter-driven sgRNA (based on the improved F+E *Spy*Cas9 scaffold), and a RSV promoter-driven GFP marker, (iii) a CMV promoter-driven AcrIIA4, AcrIIC1 or AcrIIC3 either with or without two miRNA binding sites in their 3’UTRs, or (iv) U1a promoter-driven *Nme*Cas9 co-encoding a U6 promoter-driven sgRNA (VEGFA or non-targeting control). 72 h post-transfection, cells were collected in 300 µl PBS and subsequently subjected to five freeze-thaw cycles by alternating between snap freezing in liquid nitrogen and a 37 °C water bath. The cell debris was removed by centrifugation at ∼18,000 xg for 10 min and the AAV-containing supernatant was stored at −20°C until use.

### Transductions, T7 assays, and TIDE sequencing

For T7 assays, HeLa and Huh-7 cells were seeded at a density of 3,000 cells per well, HEK293T cells were seeded at a density of 3,500 cells per well, and HL-1 cells were seeded at a density of 12,000 cells per well into 96-well plates (Eppendorf) using 100 µl culture medium per well. 16 h post-seeding, cells were co-transduced with AAV lysates encoding Cas9, a sgRNA, and the respective anti-CRISPR variant. For experiments targeting the EMX1, CCR5 or AAVS1 locus, Huh-7 and HeLa cells were transduced with 33 µl of *Spy*Cas9 vector lysate, 33 µl of EMX1, CCR5 or AAVS1 sgRNA vector lysate, and either 1, 2.5, 5, 7.5 or 10 µl of either AcrIIA4*-scaffold* or AcrIIA4*-2xmiR-122* vector lysate. For experiments targeting the VEGFA locus, Huh-7 and HEK293T cells were transduced with 40 µl of the *Nme*Cas9 AAV lysate and either 5, 10, or 20 µl of AcrIIC1 AAV lysate or 2.5, 5, or 10 µl of AcrIIC3 AAV lysate. HL-1 cells were co-transduced with AAV lysates comprising 10 µl of the *Spy*Cas9 vector, 10 µl of the Rosa-26 sgRNA vector, and 3 µl of either AcrIIA4*-scaffold* or AcrIIA4*-2xmiR-122* vector.

The volume of the AAV lysate used per well was adjusted with PBS to 100 µl for *Spy*Cas9 experiments in HeLa and Huh-7 cells, to 80 µl for *Nme*Cas9 experiments in Huh-7 and HEK293T cells, and to 23 µl for T7 assays in HL-1 cells, so that the total volume per well was identical in all samples, including the positive (Cas9 plus sgRNA, but no Acr) and negative (Cas9 only) controls. Medium was replaced 24 h (for HEK293T, HeLa and Huh-7 cells) or 4 h (for HL-1) post-infection and the transduction was repeated 24 h after the first transduction started.

72 h post- (first) transduction, cells were lysed using DirectPCR lysis reagent supplemented with proteinase K (Sigma-Aldrich). The genomic CRISPR/Cas9 target locus was PCR-amplified with primers flanking the target site (Supplementary Table 3) using Q5 Hot Start High-Fidelity DNA Polymerase (New England Biolabs). For TIDE sequencing analysis, the amplicon was purified using gel electrophoresis followed by gel extraction using the QIAquick Gel Extraction Kit (Qiagen) and by Sanger sequencing (Eurofins). Data analysis was performed using the TIDE web tool (https://tide.deskgen.com/) (37). To assess the indel frequency by T7 assay, we employed a rapid T7 protocol (26). Ten µl of the target locus amplicons were diluted 1:4 in 1x buffer 2 (New England Biolabs), heated up to 95 °C, and slowly cooled down to allow re-annealing and formation of hetero-duplexes using a nexus GSX1 Mastercycler (Eppendorf) and the following program: 95° C/5 min, 95-85° C at −2 °C per second, 85-25 °C at −0.1 °C per second. Subsequently, 0.5 µl T7 endonuclease (New England Biolabs) was added, samples were mixed and incubated at 37 °C for 15 min followed by analysis on a 2 % Tris-borate-EDTA agarose gel. The Gel iX20 system equipped with a 2.8 megapixel/14 bit scientific-grade CCD camera (INTAS) was used for gel documentation. To calculate the indel percentages from the gel images, the background was subtracted from each lane and T7 bands were quantified using the ImageJ (http://imagej.nih.gov/ij/) gel analysis tool. Peak areas were measured and percentages of insertions and deletions (indel(%)) were calculated using the formula indel (%)=100x(1-(1-fraction cleaved)^1/2^), whereas the fraction cleaved=∑(Cleavage product bands)/∑(Cleavage product bands+PCR input band). Full-length T7 assay gel images are shown in Supplementary Figure 2.

### Statistical analysis

Independent experiments correspond to cell samples seeded, transfected, and treated independently and on different days. Uncertainties in the reported mean values are indicated as the standard error of the mean (s.e.m.). A two-sided Student’s *t-*test was applied to test differences between sample groups for statistical significance. Resulting *P*-values < 0.05, 0.01 and 0.001 are indicated by one, two or three asterisks, respectively.

## RESULTS

### Design of miRNA-dependent anti-CRISPR vectors

To generate a cell-specific Cas9-ON switch, we aimed at rendering the activity of CRISPR/Cas9 dependent on the presence of cell-specific miRNAs, i.e. miRNAs that are abundant solely within the target cell type. Translating the abundance of a miRNA, which typically is a negative stimulus (causing gene expression knockdown), into a positive output (CRISPR activation) requires a negative feedback.

We hypothesized that anti-CRISPR proteins, a recently discovered class of phage-derived CRISPR/Cas inhibitors (28,30,38,39), would be an ideal mediator to establish this negative feedback. Due to their small size (∼80-150 amino acids), Acrs can be expressed quickly and efficiently from plasmids or viral vectors. More importantly, anti-CRISPR proteins block CRISPR/Cas9 DNA targeting, Cas9 nuclease function or both by directly binding to the Cas9/sgRNA complex. This post-translational inhibitory mechanism enables a complete shutdown of CRISPR/Cas9 activity even upon simultaneous delivery of Cas9, a sgRNA, and an anti-CRISPR-encoding vector (29-31,40). Coupling the expression of anti-CRISPR proteins to selected miRNAs could therefore be a suitable approach for an efficient, cell type-specific Cas9-ON switch (Figure 1A).

To validate this concept, we created a modular vector encoding a CMV promoter-driven AcrIIA4, an anti-CRISPR protein efficiently inhibiting the most widely employed Cas9 orthologue from *S. pyogenes* (30). BsmBI (Esp3l) sites present in the 3’UTR of the AcrIIA4 gene enable the introduction of miRNA target sites via Golden Gate cloning, so that AcrIIA4 expression can be set under control of any abundant, cell-specific miRNA (or set of miRNAs, see discussion).

A prominent example is miR-122, which is highly expressed in the liver, but not in any other tissue (18). Using a luciferase reporter knockdown assay, we could confirm a strong miR-122 expression in human hepatocellular carcinoma cells (Huh-7), while human cervix carcinoma (HeLa) or embryonic kidney (HEK293T) control cells showed comparably low miR-122 levels (Supplementary Figure 3).

To place AcrIIA4 under miR-122 regulation, we inserted a concatemer of two miR-122 target sites into our modular AcrIIA4 construct. We further added an N-terminal mCherry to enable fluorescence-based detection of AcrIIA4 expression (Figure 1B). Then, we co-transfected the resulting vector (mCherry-AcrIIA4*-2xmiR-122*) or a control vector lacking the miR-122 target site (mCherry-AcrIIA4*-scaffold*) alongside a *Spy*Cas9-GFP vector into Huh-7 and HeLa cells. Live-cell fluorescence microscopy and complementary Western blot analysis revealed an efficient knockdown of mCherry-AcrIIA4*-2xmiR-122* expression in Huh-7, but not in HeLa cells, thereby indicating a successful coupling of miRNA-122 abundance to AcrIIA4 expression (Figure 1C, D). As expected, *Spy*Cas9-GFP expression was not affected by the AcrIIA4 knockdown (Figure 1C, D).

### Hepatocyte- and myocyte-specific activation of *Spy*Cas9

To investigate whether the observed, miR-122-dependent knockdown of AcrIIA4 would be sufficient to release CRISPR/Cas9 activity specifically in hepatocytes, we performed a luciferase reporter cleavage assay. We co-transfected Huh-7 or HeLa cells with vectors encoding *Spy*Cas9, AcrIIA4*-2xmiR-122* (or AcrIIA4*-scaffold* as control) as well as a Firefly luciferase reporter plasmid, co-encoding a sgRNA targeting the Firefly luciferase gene, and measured luciferase activity 48 h post-transfection. As expected, we observed a significant, miR-122-dependent release of *Spy*Cas9 activity as indicated by the potent knockdown of Firefly luciferase activity observed specifically in the Huh-7 samples expressing AcrIIA4*-2xmiR-122*, but not in the AcrIIA4*-scaffold* control samples (Figure 2A). In contrast, *Spy*Cas9 was strongly inhibited in the HeLa control samples irrespective of the presence of miR-122 target sites in the AcrIIA4 3’UTR (Figure 2A), thereby confirming the functionality of our Cas9-ON switch. As expected, the degree of miR-dependent *Spy*Cas9 release in Huh-7 cells as well as *Spy*Cas9 inhibition in the off-target cells (HeLa) depended on the transfected AcrIIA4 vector dose (Figure 2A) and could be further tuned by varying the strength of the AcrIIA4-driving promoter (Supplementary Figure 4).

Next, we tested whether our Cas9-ON strategy would also enable cell type-specific editing of endogenous genomic loci. To deliver the different components of our system efficiently, we chose to employ Adeno-associated virus (AAV) vectors. AAVs are highly efficient, safe (AAVs are non-pathogenic in humans), and—very importantly—one can re-target AAVs to specific cell types or tissues by modifying the viral capsid (41–43). These properties render AAV a prime vector candidate for therapeutic gene delivery.

We packaged (i) *Spy*Cas9, (ii) sgRNAs targeting the human EMX1, CCR5 or AAVS1 locus as well as (iii) AcrIIA4*-2xmiR-122* or AcrIIA4*-scaffold* into AAV2, a well-studied AAV serotype known for its ability to efficiently transduce various cell lines (42). We then co-transduced Huh-7 or HeLa cells with combinations of these vectors, while varying the AcrIIA4 vector dose, and measured the frequency of insertions and deletions at the target loci using a T7 endonuclease assay and TIDE sequencing (37). We observed potent, miRNA-122-dependent gene editing at all three target loci in Huh-7 cells with a dynamic range of regulation of up to 16-fold (Figure 2B-D, Supplementary Figure 5). The editing efficiency and leakiness of the system in the OFF-state depended on the used AcrIIA4 vector dose (Figure 2B-D). Importantly, in the HeLa control cells, Cas9 activity was equally suppressed in the presence of the AcrIIA4*-2xmiR-122* or AcrIIA4*-scaffold* vector (Figure 2B), indicating that our miRNA-122-dependent Cas9-ON switch is indeed hepatocyte-specific.

The large size of CRISPR/*Spy*Cas9 (∼ 1,300 amino acids) poses a challenge with respect to its efficient delivery and expression, in particular when using vectors with a constrained packaging capacity. To circumvent this problem, several groups have recently developed split-*Spy*Cas9 variants, which comprise an N-and C-terminal Cas9 fragment that reconstitute a functional Cas9 when co-expressed within the same cell (44–47). To test whether our anti-CRISPR-based Cas9-ON strategy would also work for split-*Spy*Cas9s, we employed an intein-based split-*Spy*Cas9 system recently reported by the lab of George Church (48). We co-transfected plasmids encoding the N- and C-terminal *Spy*Cas9 fragments alongside an AcrIIA4 vector (with or without miR-122 sites in the 3’UTR) and the aforementioned luciferase cleavage reporter into Huh-7 cells (or HeLa cells as control). In Huh-7 cells, the split-*Spy*Cas9 remained fully active in the presence of the AcrIIA4*-2xmiR-122* vector as indicated by potent luciferase knockdown, but was completely impaired if we co-administered the AcrIIA4*-scaffold* construct (Supplementary Figure 6). In HeLa cells, in contrast, split-*Spy*Cas9 activity was impaired upon co-delivery of both, the AcrIIA4*-2xmiR-122* or the AcrIIA4*-scaffold* vector (Supplementary Figure 6), demonstrating that our Cas9-ON switch can also be applied to control split-*Spy*Cas9.

MiR-1 plays an important role in muscle cell differentiation (22,23) and remains highly expressed in mature muscle cells (49). Using a luciferase knockdown assay, we confirmed high miR-1 levels in HL-1 cells, a widely employed murine cardiac muscle cell model (Supplementary Figure 7). We hypothesized that, similarly to miR-122 in hepatocytes, miR-1 could be harnessed for myocyte-specific activation of CRISPR/Cas9 using the identical, Acr-based strategy. We therefore exchanged the two miR-122 binding sites in our AcrIIA4 constructs by two miR-1 target sites (Figure 2E). Then, we packaged the resulting AcrIIA4*-2xmiR-1* construct or the AcrIIA4*-scaffold* construct (as control), a *Spy*Cas9 transgene, and a sgRNA targeting the murine Rosa-26 locus into AAV serotype 6, which is known to efficiently transduce a wide spectrum of tissues *in vitro and vivo*, including myocytes (50–52). Upon co-infection of HL-1 cells with these vectors, we observed potent editing of the Rosa-26 locus in the presence of the AcrIIA4-*2xmiR-1* vector, but not when using the AcrIIA4-*scaffold* control vector (Figure 2F,G and Supplementary Figure 8), thereby demonstrating successful miR-1-dependent release of *Spy*Cas9 activity in myocytes.

### miRNA control of dCas9-effector fusions

So far, we have demonstrated the power of our Cas9-ON system for cell type-specific genome editing. However, the CRISPR/Cas9 system offers many applications that go beyond a targeted introduction of double-strand breaks. These are typically based on catalytically inactive d(ead)Cas9 mutants employed as programmable DNA binding domain to recruit effector domains to selected genomic loci. These effectors then mediate e.g. transcriptional activation or repression (6–8,53–55), epigenetic modification (9–11,56,57) or base editing (12,13,58). AcrIIA4 inhibits the *Spy*Cas9 mainly by blocking its DNA binding (31,59). We therefore hypothesized that our miRNA-dependent *Spy*Cas9-ON switch should also be applicable to *Spy*dCas9-effector fusions.

To test this hypothesis, we co-transfected vectors encoding a *Spy*dCas9-VP64 transcriptional activator, a Tet operator (TetO) targeting sgRNA, a luciferase reporter driven from a TetO-dependent promoter, and an AcrIIA4*-2xmiR-122* or AcrIIA4*-scaffold* vector into Huh-7 cells (Figure 3A). We observed a potent, miR-122-dependent release of luciferase reporter expression with a dynamic range of regulation of up to 114-fold (Figure 3B). Remarkably, the leakiness and dynamic range of the system could be tuned over a wide range by varying the AcrIIA4 vector dose (Figure 3B). These results illustrate that our *Spy*Cas9-ON system can also be applied for cell type-specific activation of *Spy*dCas9-effector fusions.

### Hepatocyte-specific activation of *Nme*Cas9

Although the *Spy*Cas9 remains the most widely employed CRISPR/Cas orthologue, the ongoing discovery and characterization of novel CRISPR/Cas effectors from various species rapidly expands the CRISPR toolbox. For many of these novel type I and II CRISPR/Cas effectors, corresponding anti-CRISPR proteins have already been found or are likely to be discovered in the near future (28–30,38,40,60,61), suggesting that our Cas9-ON approach might be easily transferable to many other CRISPR/Cas orthologues. One such orthologue is the *Neisseria meningitidis* (*Nme)*Cas9, which is not only ∼ 300 amino acids smaller than *Spy*Cas9, but also shows a far lower activity on off-target loci, presumably due to its extended protospacer sequence (62–64). Two anti-CRISPR proteins have recently been described, which efficiently inhibit *Nme*Cas9 via distinct mechanisms. AcrIIC1 perturbs the *Nme*Cas9 nuclease function, while AcrIIC3 induces *Nme*Cas9 dimerization, thereby impairing its DNA binding (29,32).

We speculated that, similar to *Spy*Cas9 control via miR-dependent AcrIIA4, placing AcrIIC1 and AcrIIC3 under miRNA regulation would enable cell-specific activation of *Nme*Cas9. To test this hypothesis, we codon-optimized the AcrIIC1 and AcrIIC3 genes, introduced miR-122 target sites into their 3’UTRs and packaged them into AAV2. Then, we co-transduced Huh-7 or HEK293T control cells with the AcrIIC1*-2xmiR-122* or AcrIIC3-*2xmiR-122* vector (or AcrIIC1*-scaffold* or AcrIIC3*-scaffold* as control) alongside a vector co-encoding *Nme*Cas9 and a sgRNA targeting the human VEGFA locus (65) (Amrani *et al*., 2018, BioRxiv: https://doi.org/10.1101/172650). As hoped for, we observed potent miR-122-dependent indel mutation of the VEGFA target locus with an up to 24-fold dynamic range of miRNA-regulation depending on the vector dose (Figure 4 and Supplementary Figure 9). In HEK293T control cells, in contrast, *Nme*Cas9 inhibition was independent of the presence of miR-122 target sites on the Acr vectors (Figure 4), demonstrating that our Cas-ON switch enables cell-specific activation of *Nme*Cas9.

## DISCUSSION

CRISPR/Cas technologies enable detailed genetic studies in cells and animals and hold unmet potential for the treatment of genetic disorders. To render CRISPR/Cas *in vivo* applications as specific and thus as safe as possible, strategies to confine the activity of Cas9 nucleases or dCas9-based effectors to defined cells and tissues are highly desired.

In this study, we employed cellular miRNA signatures to control the expression of anti-CRISPR proteins, thereby creating a synthetic circuit efficiently confining Cas9 activity to selected target cells. Liver and muscle are interesting target tissues for CRISPR-mediated gene therapy approaches, e.g. for treatment of hemophilia or Duchenne muscular dystrophy, respectively (66–70). Using our Cas9-ON system, we were able to activate CRISPR/Cas9 selectively in hepatocytes or myocytes by employing miR-122 and miR-1 as cellular markers, respectively, while efficiently suppressing Cas9 in unrelated cell types lacking these markers (HeLa and HEK293T). As specific miRNAs or miRNA signatures have been discovered for a variety of cell types (18,19), e.g. hematopoietic cells (miR-142 (24,71,72)) or neurons (e.g. miR-376a, miR-434 (73)), we anticipate that our Cas9-ON strategy should be highly useful in the context of various applications. Incorporating target sites for multiple, abundant miRNAs into the Acr gene 3’UTR and/or coupling our Cas9-ON switch with existing, miRNA-based Cas9-OFF switches (26,27) might even further improve the level of specificity that can be achieved with our system.

In any case, the Acr vector dose is a crucial parameter to consider. We showed that too high a dose of Acr vector may cause significant suppression of CRISPR/Cas9 activity, even within the target cell population, presumably due to a limited capacity of the RNAi machinery. In contrast, too low a dose of Acr vector leads to insufficient Cas9 inhibition in off-target cells. Importantly, by modulating the dose of the supplied Acr vector or the strength of the Acr-driving promoter, one can easily tune the system toward the optimal switching behaviour for a particular application.

Another important feature of our Cas9-ON switch is its modularity, i.e. it should be compatible with any Cas9 orthologue, for which a potent anti-CRISPR protein is known (as exemplified here for *Spy*Cas9 and *Nme*Cas9). In light of the ongoing, rapid discovery and characterization of novel Acrs, the application spectrum of our switch is likely to further expand in the near future. Importantly, our Cas-ON strategy is also applicable to dCas9-effector fusions, provided an Acr is employed, which impairs Cas9 DNA binding. This is the case, e.g. for AcrIIA4 and AcrIIC3, which block DNA targeting of *Spy*Cas9 and *Nme*Cas9, respectively, but not for AcrIIC1, which impairs the *Nme*Cas9 nuclease function, but does not interfere with its DNA binding. Therefore, the underlying, inhibitory mechanism can be an important parameter to consider when selecting Acrs to be used in our Cas-ON system.

Importantly, our Cas-ON is compatible with many existing strategies for tissue-specific gene delivery and expression, such as engineered or evolved AAV vectors (41,42,74) or tissue-specific promoters (75,76). Thus, combining these approaches, potentially with additional layers of CRISPR/Cas control via chemical triggers (44,77,78) or light (34,46,79,80), will likely enable highly specific genome perturbations.

While we foresee that the most relevant applications of our approach will be in animal models and, in the long run, potentially in human patients, we reckon that a careful investigation of toxicity or immune reactions that might result from Acr overexpression should precede *in vivo* translation of our Cas9-ON strategy. Moreover, to avoid continuous sequestration of the endogenous miRNA pool within the target cells, it could be advisable to couple our Cas9-ON strategy to vector self-inactivation (81). Taken together, our work demonstrates the power of miRNA-dependent anti-CRISPR transgenes to confine CRISPR/Cas9 activity to selected cells types and facilitate safe and precise genome perturbations in animals and patients.

## Supporting information

## AVAILABILITY

Vectors will be made available via Addgene (#120293 - 120303).

## SUPPLEMENTARY DATA

Supplementary Figures 1-9 and Supplementary Tables 1-3 are provided as Supplementary Information.

## ACKNOWLEDGEMENT

We thank the Synthetic Biology group (Heidelberg University), the Virus-Host Interactions group (University Heidelberg, Hospital) and the Intelligent Imaging group (DKFZ, Heidelberg) for helpful discussions, K. Rippe (DKFZ, Heidelberg) and E. Wiedtke (University Hospital, Heidelberg) for providing cell lines, J. Notbohm and A. Bietz (both Universitiy Heidelberg) for testing the VEGFA sgRNA and K. Börner (University Hospital, Heidelberg) for sharing expertise on AAVs. We are further grateful to Moritoshi Sato (University of Tokyo), Feng Zhang (Massachusetts Institute of Technology, Cambridge), George Church (Harvard University, Cambridge), and Erik J. Sontheimer (University of Massachusetts Medical School, Worcester) for sharing plasmids.

## Author contributions

D.G. and D.N. conceived the study. M.D.H. and D.N. designed experiments. M.D.H., S.A., S.G., K.R., and M.M. conducted experiments. M.D.H. and D.N. analysed and interpreted data. J.F. developed the luciferase cleavage reporter construct. D.G. provided expertise for AAV experiments and CRISPR assays. C.D. made practical contributions. R.E., D.G., and D.N. jointly directed the work. M.D.H., D.G., and D.N. wrote the manuscript with support from all authors.

## FUNDING

This study was funded by the Helmholtz association, the German Research Council (DFG) and the Federal Ministry of Education and Research (BMBF). M.D.H. was supported by a Helmholtz International Graduate School for Cancer Research scholarship (DKFZ, Heidelberg). D.G. is grateful for funding from the German Center for Infection Research (DZIF) [grant number TTU-HIV 04.803]. J.F. and D.G. were supported by the Cystic Fibrosis Foundation (CFF) [grant number GRIMM15XX0]. D.G. and R.E. acknowledge funding from the Transregional Collaborative Research Center TRR179 (DFG; TP18 to D.G. and TPX to R.E.). J.F. and D.G. acknowledge additional funding by the Collaborative Research Center SFB1129 (TP2; 2014-2018) and the Cluster of Excellence CellNetworks (DFG) [grant number EXC81]. D.G. and C.D. participate in the SMART-HaemoCare consortium, which receives financial support from ERA-NET E-Rare-3 [JCT-2015) / DFG]. K.R. was partially supported by the Marie Curie International Incoming Fellowship [PIIF-GA-2013-627329].

## CONFLICT OF INTEREST

D.N., D.G., R.E., M.D.H., J.F., C.D. and S.A. have filed patent applications related to this work.

