## Supplementary material for "Cell-specific CRISPR/Cas9 activation by microRNA-dependent expression of anti-CRISPR proteins"

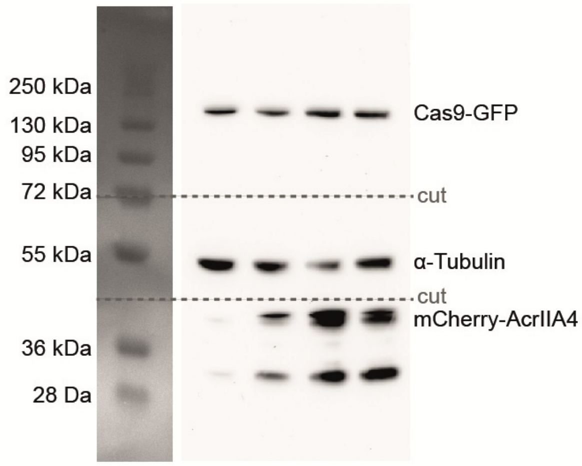


**Supplementary Figure 1.** Full-length Western blot image (corresponds to Figure 1D). The ladder is the PageRuler Prestained Protein Ladder (ThermoFisher). Positions at which the membrane was cut prior to antibody incubation are indicated (dashed lines).

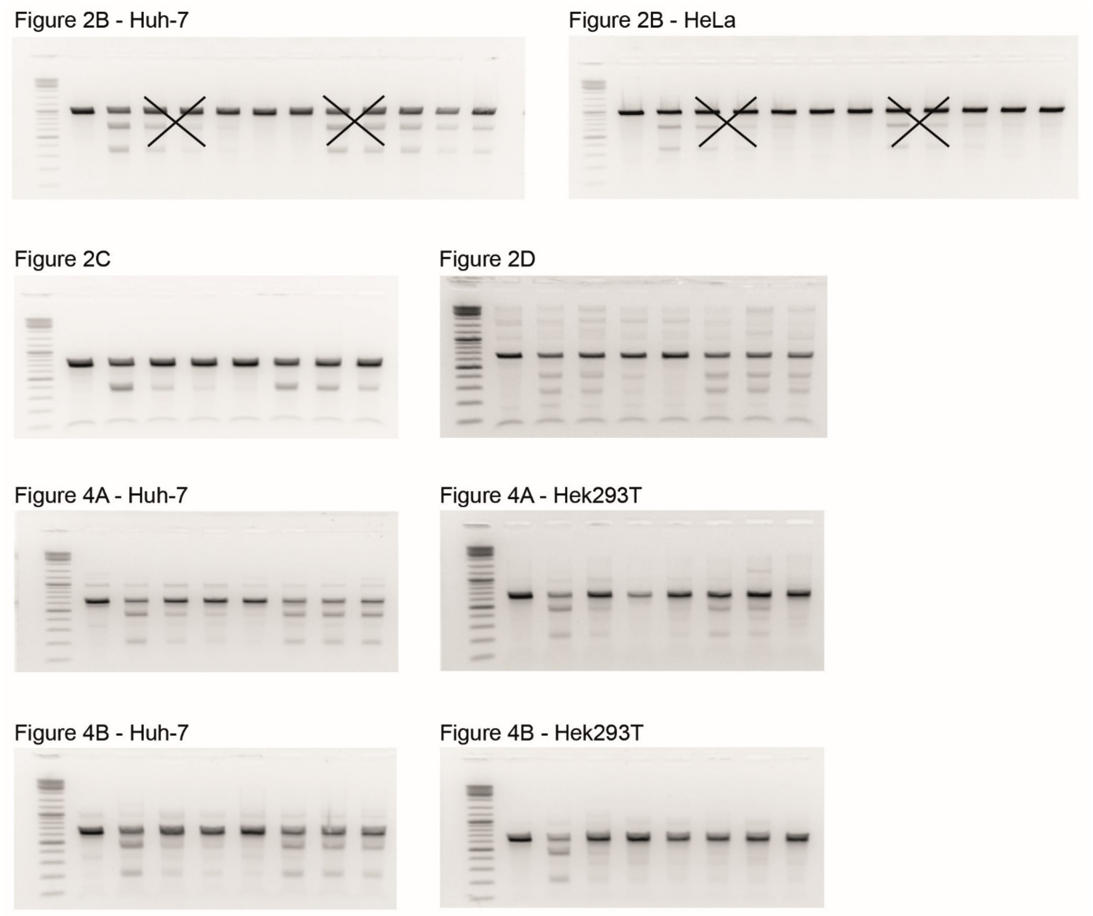


**Supplementary Figure 2.** Full-length gel images of T7 endonuclease assays. The ladder is the Gene Ruler DNA Ladder Mix (Thermo Fisher).

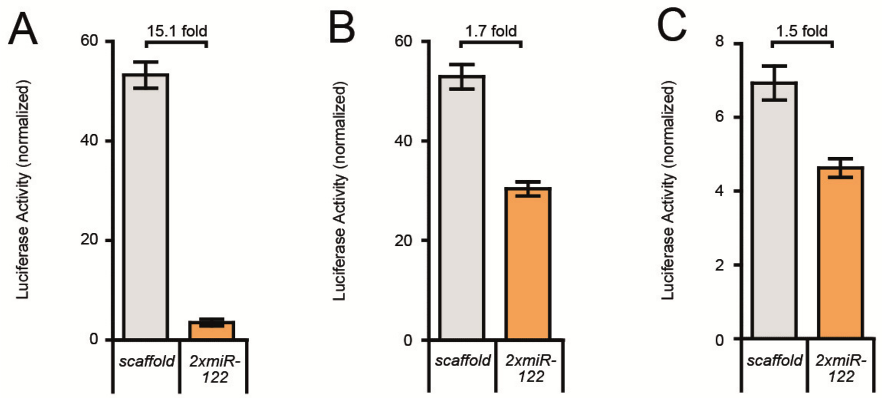


**Supplementary Figure 3.** MiRNA-122 is highly abundant specifically in hepatocytes. Huh-7 (**A**), HeLa (**B**), and HEK293T (**C**) cells were transfected with a luciferase reporter construct carrying two miR-122 binding sites in the 3'UTR (2xmiR-122) or not (scaffold, as control), followed by luciferase assay. The reporter bearing the miR-122 binding sites is efficiently knocked down in Huh-7 cells, but only mildly affected in HeLa or HEK293T cells. Data are means ± s.e.m. (n = 3 independent experiments).

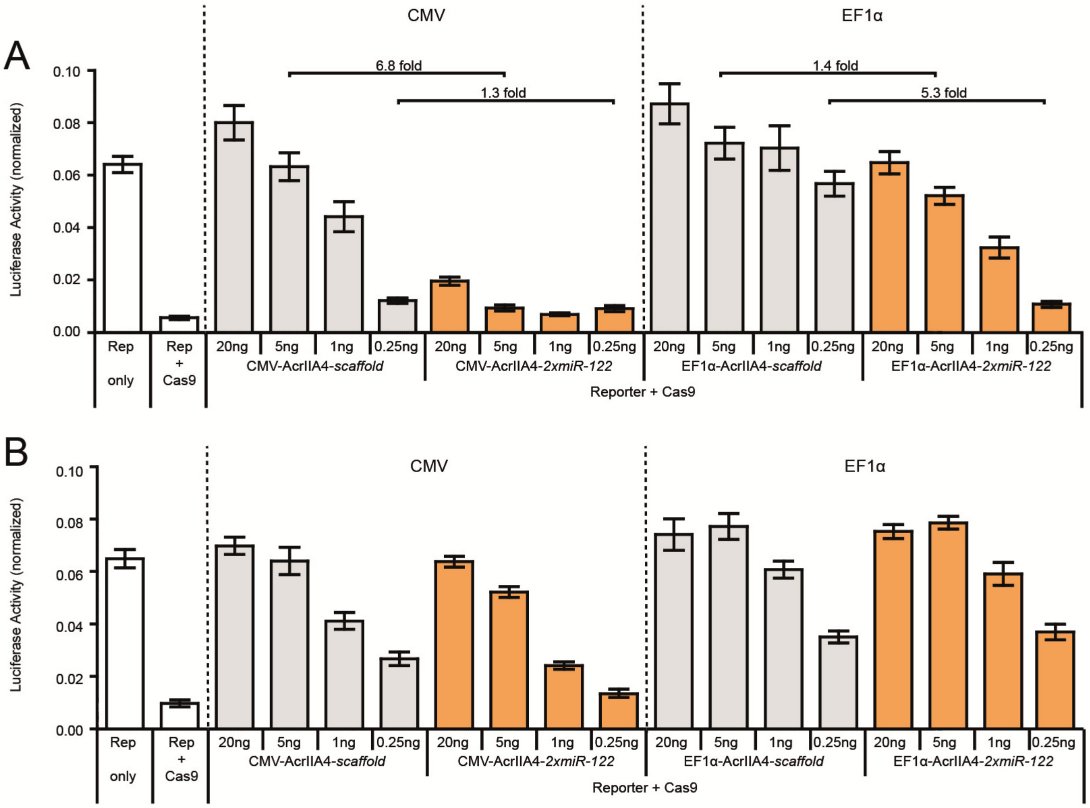


**Supplementary Figure 4.** *Spy*Cas9 inhibition can be modulated by tuning the strength of the AcrIIA4-driving promoter. Huh-7 cells (**A**) and HeLa (**B**) cells were co-transfected with constructs encoding *Spy*Cas9, a luciferase reporter, a reporter-targeting sgRNA, and AcrIIA4-*2xmiR-122* or AcrIIA4-*scaffold* as control, followed by luciferase assay. A CMV promoter or an EF1α promoter was used to drive Acr expression and the Acr vector dose was varied during transfection as indicated. Data are means ± s.e.m. (n = 3 independent experiments).

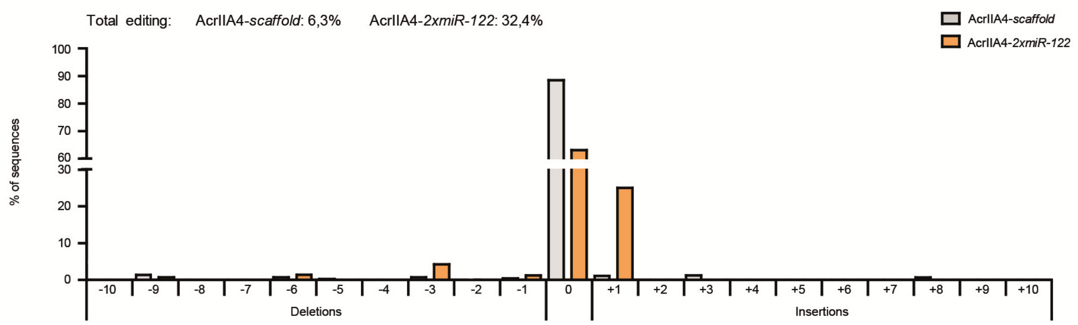


**Supplementary Figure 5.** MiR-122-dependent editing of the EMX1 locus in hepatocytes. Huh-7 cells were co-transduced with AAV vectors expressing *Spy*Cas9, a sgRNA targeting the human EMX1 locus, and AcrIIA4-*2xmiR-122* or AcrIIA4-*scaffold* (as control). TIDE sequencing revealed a high frequency of insertions and deletions in the AcrIIA4-*2xmiR-122*, but not in the AcrIIA4-*scaffold* sample. The total editing efficiencies as calculated by the TIDE algorithm are indicated on top. Data for a representative sample is shown.

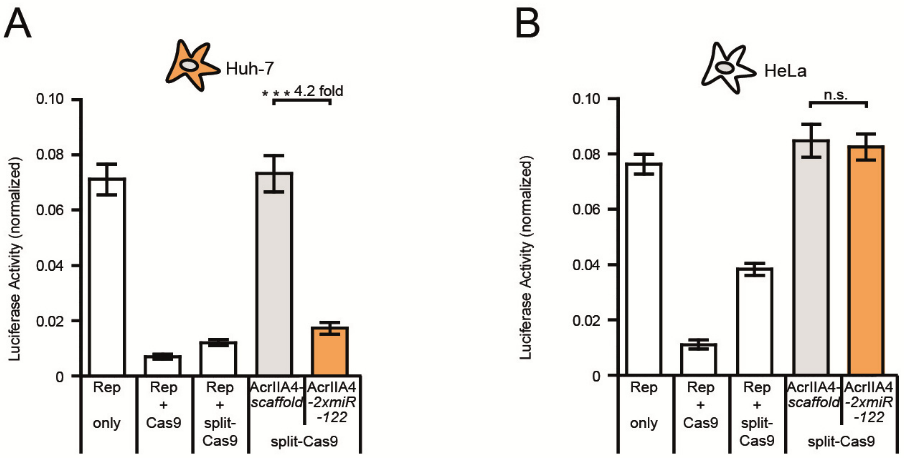


**Supplementary Figure 6.** Hepatocyte-specific activation of split-*Spy*Cas9. Huh-7 (**A**) and HeLa (**B**) cells were co-transfected with constructs expressing a luciferase reporter, a reporter-targeting sgRNA, an N- and C-terminal *Spy*Cas9 fragment fused to split-inteins, and AcrIIA4-*2xmiR-122* or AcrIIA4-*scaffold* (as control), followed by luciferase assay. Data are means ± s.e.m. (n = 4 independent experiments). n.s. = not significant, ***P < 0.001, by two-sided Student’s *t*-test.

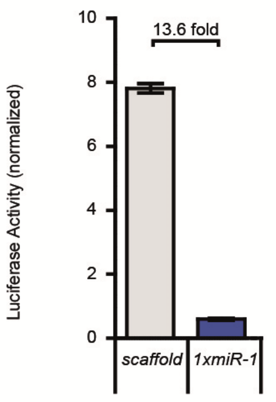


**Supplementary Figure 7.** MiRNA-1-mediated reporter knockdown in murine cardiac myocytes. HL-1 cells were transduced with AAVs encoding a luciferase reporter construct carrying a miR-1 target site in the 3’UTR (1xmiR-1) or not (scaffold, as control), followed by luciferase assay. Data are means ± s.e.m. (n = 3 replicates, i.e. parallel transfections).

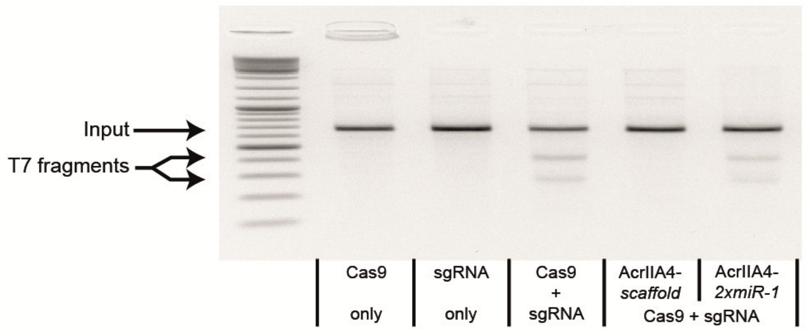


**Supplementary Figure 8.** MiR-1-dependent gene editing in myocytes. HL-1 cells were co-transduced with AAV vectors encoding *Spy*Cas9, a sgRNA targeting the Rosa-26 locus, and either AcrIIA4-*2xmiR-1* or AcrA4-*scaffold* (control), followed by T7 endonuclease assay. Data for a representative sample is shown.

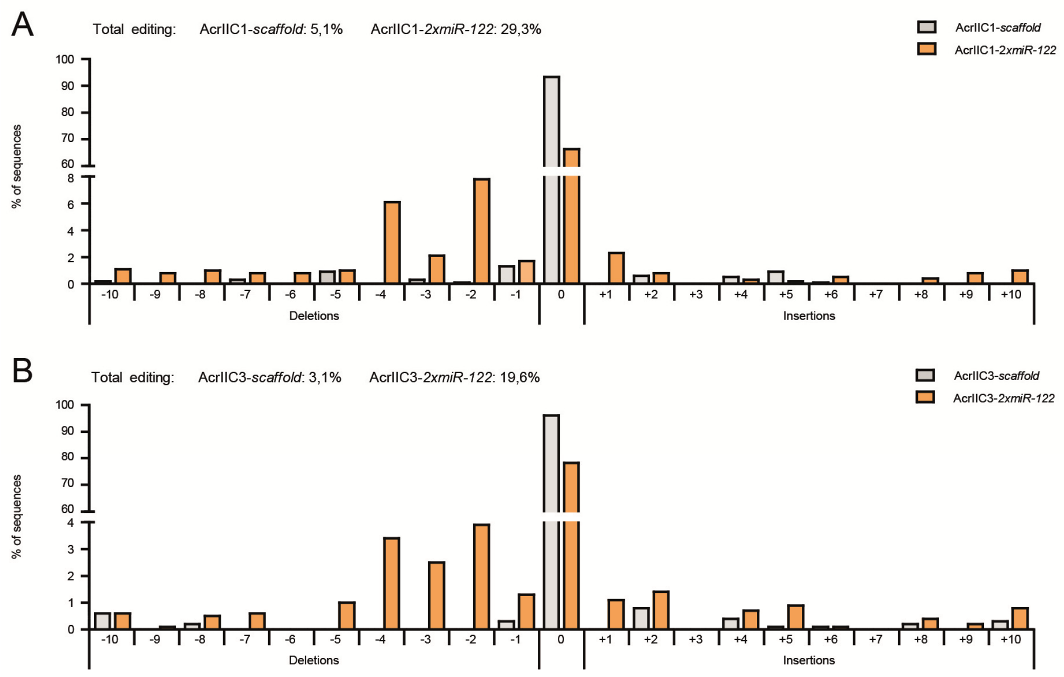


**Supplementary Figure 9.** MiR-122-mediated *Nme*Cas9 activation. Huh-7 cells were co-transduced with AAV vectors encoding *Nme*Cas9, a sgRNA targeting the human VEGFA locus, and either AcrIIC1-*2xmiR-122* or AcrIIC1-*scaffold* (**A**), or AcrIIC3-*2xmiR-122* or AcrIIC3-*scaffold* (**B**). TIDE sequencing revealed a broad range of insertions and deletions for the AcrIIC1-*2xmiR-122*/AcrIIC3-*2xmiR-122* samples, but not for the AcrIIC1-*scaffold*/AcrIIC3-*scaffold* control samples. The total editing efficiencies as calculated by the TIDE algorithm are indicated on top. Data for a representative samples are shown.

**Supplementary Table 1.** List of constructs created and used in this study. ITR, inverted terminal repeat. TK, thymidine kinase. SV40, simian virus 40. BGH, bovine growth hormone. LTR, long terminal repeat. CMV, cytomegalovirus. RSV, respiratory syncytial virus. EF1α, elongation factor 1α

| No | Name/Group | Insert description |
|---|---|---|
| | **Reporter** | |
| **1** | Luciferase cleavage reporter (to measure *Spy*Cas9 activity) | 3'ITR, SV40 promoter, *Renilla luciferase*, TK promoter, Firefly luciferase, SV40 polyA, H1 promoter, sgRNA(Firefly luciferase), 5' ITR |
| **2** | Tet-inducible luciferase reporter (Addgene # 64127, to measure *Spy*dCas9-VP64 activity) | Tet-responsive elements, ECFP, CMV promoter, Firefly luciferase, SV40 polyA |
| **3** | pRL-TK (Promega) | TK promoter, *Renilla* luciferase, SV40 polyA |
| **4** | pSiCheck-2 (no miR target site, control) | SV40 promoter, *Renilla* luciferase, scaffold, BGH polyA, TK promoter, Firefly luciferase |
| **5** | pSiCheck-2 (2xmiR-122 target sites) | SV40 promoter, *Renilla* luciferase, 2xmiR-122 binding sites, BGH polyA, TK promoter, Firefly luciferase |
| **6** | pAAV-pSi (no miR target site control) | 3'ITR, SV40 promoter, *Renilla* luciferase, TK promoter, Firefly luciferase, SV40 promoter, 5'ITR |
| **7** | pAAV-pSi (1xmiR-1 target site) | 3'ITR, SV40 promoter, *Renilla* luciferase, 1xmiRNA-1 binding site, TK promoter, Firefly luciferase, SV40 polyA, 5'ITR |
| | **Cas9 Variants** | |
| **8** | CMV-*Spy*Cas9 (Addgene # 113033) | CMV promoter, NLS-*Spy*Cas9-NLS, BGH polyA |
| **9** | *Spy*Cas9-GFP | CMV promoter, NLS-*Spy*Cas9-NLS-GFP, BGH polyA |
| **10** | dCAS9-VP64_GFP (Addgene # 61422) | CMV promoter, 5'LTR(HIV-1), EF1α promoter, NLS-*Spy*dCas9-NLS-VP64(Transactivator)-T2A-EGFP, 3'LTR(HIV-1) |
| **11** | pAAV-CMV-*Spy*Cas9 (Addgene # 113034) | 3'ITR, minimal CMV promoter, NLS-*Spy*Cas9-NLS, 5'ITR |
| **12** | AAV N-*Spy*Cas9-Intein, U6-sgRNA scaffold (F+E) (Addgene # 120293) | 3'ITR, SMVP promoter (Reference 1), NLS-*Spy*Cas9(N)-N-intein, SV40 polyA, U6, *Spy*Cas9 sgRNA scaffold(F+E), 5'ITR |
| **13** | AAV Intein-C-*Spy*Cas9 | 3'ITR, SMVP promoter, C-intein-*Spy*Cas9(C), SV40 polyA, XbaI site, MluI site, 5'ITR |
| **14** | AAV Intein-C-*Spy*Cas9, CMV-AcrIIA4-*scaffold* (2xBsmBI sites) (Addgene # 120294) | 3'ITR, SMVP promoter, C-intein-*Spy*Cas9(C), SV40 polyA, CMV promoter, AcrIIA4, scaffold sequence, BGH polyA, 5'ITR |
| **15** | AAV Intein-C-*Spy*Cas9, CMV-AcrIIA4-*2xmiR-122* target sites (Addgene # 120295) | 3'ITR, SMVP promoter, C-intein-*Spy*Cas9(C), SV40 polyA, CMV promoter, AcrIIA4, 2xmiR-122 binding sites, BGH polyA, 5'ITR |
| **16** | pEJS654 All-in-One AAV-sgRNA-h*Nme*Cas9 (Addgene # 112139) | AAV2-ITR, *Nme*Cas9 sgRNA scaffold, U6 promoter, U1a promoter, NLS-*Nme*Cas9-NLSs, β-globin polyA, AAV2-ITR |
| **17** | All-in-One AAV-sgRNA-VEGFA-h*Nme*Cas9 | AAV2-ITR, *Nme*Cas9-sgRNA(VEGFA), U6 promoter, U1a promoter, NLS-*Nme*Cas9-NLSs, β-globin polyA, AAV2-ITR |

Supplementary Table 1 continued

**sgRNAs**

| | | |
|---|---|---|
| **18** | sgRNA1_Tet-inducible Luciferase reporter (Addgene # 64161) | U6 promoter, *Spy*Cas9-sgRNA(Tet-inducible promoter) |
| **19** | pAAV-RSV-GFP-U6-EMX1 sgRNA (*Spy*Cas9 scaffold) (Addgene # 113040) | AAV2-ITR, RSV promoter, EGFP, U6 promoter, *Spy*Cas9-sgRNA(EMX1), AAV4-ITR |
| **20** | pAAV-RSV-GFP-U6-Rosa-26 sgRNA (*Spy*Cas9 scaffold) (Addgene # 120296) | AAV2-ITR, RSV promoter, EGFP, U6 promoter, *Spy*Cas9-sgRNA(Rosa-26), AAV4-ITR |
| **21** | pAAV-RSV-GFP-U6-CCR5 sgRNA (*Spy*Cas9 scaffold) (Addgene # 113041) | AAV2-ITR, RSV promoter, EGFP, U6 promoter, *Spy*Cas9-sgRNA(CCR5), AAV4-ITR |
| **22** | pAAV-RSV-GFP-U6-AAVS1 sgRNA (*Spy*Cas9 scaffold) | AAV2-ITR, RSV promoter, EGFP, U6 promoter, *Spy*Cas9-sgRNA(AAVS1), AAV4-ITR |

**Anti-CRISPR Variants**

| | | |
|---|---|---|
| **23** | CMV-mCherry-AcrIIA4-*scaffold* | CMV promoter, NLS-mCherry-AcrIIA4, scaffold sequence, BGH polyA |
| **24** | CMV-mCherry-AcrIIA4 –2x*miR-122* | CMV promoter, NLS-mCherry-AcrIIA4, 2xmiR-122 binding sites, BGH polyA |
| **25** | CMV-AcrIIA4-*scaffold* | CMV promoter, AcrIIA4, scaffold sequence, BGH polyA |
| **26** | CMV-AcrIIA4-2x*miR-122* | CMV promoter, AcrIIA4, 2xmiR-122 binding sites, BGH polyA |
| **27** | AAV EF1α-AcrIIA4-*scaffold* | AAV4-ITR, H1 promoter, *Spy*Cas9-sgRNA-scaffold, EF1α promoter, AcrIIA4, scaffold sequence, BGH polyA, AAV2-ITR |
| **28** | AAV EF1α- AcrIIA4-2x*miR-122* | AAV4-ITR, H1 promoter, *Spy*Cas9-sgRNA-scaffold, EF1α promoter, AcrIIA4, 2xmiR-122 binding sites, BGH polyA, AAV2-ITR |
| **29** | AAV CMV-driven AcrIIA4-*scaffold* (Addgene # 120297) | AAV4-ITR, U6 promoter, *Spy*Cas9-sgRNA-scaffold, CMV promoter, AcrIIA4, scaffold sequence, BGH polyA, AAV2-ITR |
| **30** | AAV CMV-driven AcrIIA4-2x*miR-122* (Addgene # 120298) | AAV4-ITR, U6 promoter, *Spy*Cas9-sgRNA-scaffold, CMV promoter, AcrIIA4, 2xmiR-122 binding sites, BGH polyA, AAV2-ITR |
| **31** | AAV CMV-driven AcrIIA4-2x*miR-1* (Addgene # 120299) | AAV4-ITR, U6 promoter, *Spy*Cas9-sgRNA-scaffold, CMV promoter, AcrIIA4, 2xmiR-1 binding sites, BGH polyA, AAV2-ITR |
| **32** | AAV CMV-driven AcrIIC1-*scaffold* (Addgene # 120300) | AAV4-ITR, U6 promoter, *Spy*Cas9-sgRNA-scaffold, CMV promoter, AcrIIC1, scaffold sequence, BGH polyA, AAV2-ITR |
| **33** | AAV CMV-driven AcrIIC3-*scaffold* (Addgene # 120301) | AAV4-ITR, U6 promoter, *Spy*Cas9-sgRNA-scaffold, CMV promoter, AcrIIC3, scaffold sequence, BGH polyA, AAV2-ITR |
| **34** | AAV CMV-driven AcrIIC1-2x*miR-122* (Addgene # 120302) | AAV4-ITR, U6 promoter, *Spy*Cas9-sgRNA-scaffold, CMV promoter, AcrIIC1, 2xmiR-122 binding sites, BGH polyA, AAV2-ITR |
| **35** | AAV CMV-driven AcrIIC3-2x*miR-122* (Addgene # 120303) | AAV4-ITR, U6 promoter, *Spy*Cas9-sgRNA-scaffold, CMV promoter, AcrIIC3, 2xmiR-122 binding sites, BGH polyA, AAV2-ITR |

Supplementary Table 1 continued

| | **AAV Production Plasmids** | |
|---|---|---|
| **36** | Adenoviral helper plasmid | Ad2 VA RNA, Ad2 E4, Ad2 E2A |
| **37** | WHc2 | AAV2 cap, AAV2 rep |
| **38** | WHc6 | AAV6 cap, AAV2 rep |

| | **Stuffer DNA** | |
|---|---|---|
| **39** | pcDNA3.1$^{(-)}$ | empty vector (inert DNA stuffer) |

**Supplementary Table 2.** sgRNA target sites. Sequences are in 5' to 3' direction; the PAM sequence is indicated in bold. References indicate the publications originally reporting the corresponding sgRNAs.

| **Gene (Reference)** | **Cas9 orthologue** | **Target Sequence** |
|---|---|---|
| Firefly luciferase | *Spy*Cas9 | GGACTCTAAGACCGACTACC**AGG** |
| Tet-responsive element (2) | *Spy*Cas9 | TCTCTATCACTGATAGGGAG**TGG** |
| EMX1 (2) | *Spy*Cas9 | GAGTCCGAGCAGAAGAAGAA**GGG** |
| Rosa-26 (3) | *Spy*Cas9 | ACTCCAGTCTTTCTAGAAGA**TGG** |
| CCR5 (2) | *Spy*Cas9 | TGACATCAATTATTATACAT**CGG** |
| AAVS1 (4) | *Spy*Cas9 | GGGCCACTAGGGACAGGAT**TGG** |
| VEGFA (Amrani *et al*., 2018, BioRxiv: https://doi.org/10.1101/172650) | *Nme*Cas9 | GCGGGGAGAAGGCCAGGGGTCACT**CCAGGATT** |

**Supplementary Table 3.** Primers used for genomic PCRs. Sequences are in 5' to 3' direction.

| **Locus (species)** | **Direction** | **Sequence** |
|---|---|---|
| EMX1 (human) | forward | GGAGCAGCTGGTCAGAGGGG |
| | reverse | GGGAAGGGGGACACTGGGGA |
| Rosa-26 (mouse) | forward | CGTGCAAGTTGAGTCCATCCGCC |
| | reverse | ACTCCGAGGCGGATCACAAGCA |
| CCR5 (human) | forward | GAGCCAAGCTCTCCATCTAGT |
| | reverse | GCCCTGTCAAGAGTTGACAC |
| AAVS1 (human) | forward | TCCAGGGGTCCGAGAGCTCAGCTAG |
| | reverse | CCAGACAGCCGCGTCAGAGCAGCTC |
| VEGFA (human) | forward | TCCAGATGGCACATTGTCAG |
| | reverse | AGGGAGCAGGAAAGTGAGGT |
